# Genome mutations and environmental perturbations reshape the epigenomic topology to drive functional collapse in digital cells

**DOI:** 10.64898/2026.08.24.746697

**Authors:** Claudia Moreno-Fernandez-Aliseda, Miguel A. Fortuna, Daniel Rico

## Abstract

Understanding how genetic and environmental perturbations drive cellular dysfunction remains a challenge. Using digital somatic cells, we evaluate the functionality of over eight thousand genomes across thousands of environments alongside systematic mutational screens. We find that environmental perturbations induce loss of function with a higher probability than point mutations. Crucially, however, most mutations causing loss of function, rather than acting as direct sequence-level damage, converge with environmental stress by triggering reorganization of the epigenomic topology. Our data suggest that loss of functionality is best understood not as cumulative damage, but as a perturbation-driven sate transition within a reconfigurable epigenetic landscape.

## Background

The function of a cell emerges from the interaction between genetic information and regulatory states shaped by the environment [1]. Loss of cellular function refers to impairment of biological activities, often associated with mitochondrial dysfunction, oxidative stress, or cellular senescence. Disruptions in cellular function are central to the pathogenesis of multiple diseases and age-associated degeneration [2], highlighting the importance of understanding the mechanisms that establish and destabilize these states.

Both genome mutations and enviromental perturbations have traditionally been implicated in driving these functional transitions. However, a major challenge in disentangling these contributions lies in the intrinsic interdependence between the genome and the epigenome [3], that is highly dynamic, reversible and strongly influenced by environmental, developmental, and aging-related signals that can ultimately lead to functional decline [4]. DNA mutations can disrupt coding or regulatory elements [5], whereas epigenetic changes can modify transcriptional states without altering DNA sequence [4]. Moreover, genetic changes can reshape epigenetic states [6], while epigenetic configurations can influence gene expression, genome stability, and even mutation rates, creating a bidirectional coupling that makes establishing causality almost intractable in natural systems [7].

We investigated how environmental and genetic perturbations affect viability, reorganize epigenomic states, and ultimately shape functional stability in digital cells. We use the Avida platform as a digital twin model of somatic cells that allows to explore how these mechanisms contribute to functional collapse. Digital organisms within Avida are self-replicating computer programs that behave like somatic cells, dividing asexually in a miotic-like manner and evolving through mutations in a controlled environment [8,9]. Recent work has highlighted the Avida system as a key model for studying the evolution of artificial intelligence programs [10]. In our context, its architecture enables precise mechanistic analyses of how digital genomes and their execution respond to perturbations.

## Results and Discussion

The genome of a digital cell consists of executable instructions whose context-dependent execution gives rise to functional phenotypes (**Figure 1A**) - Boolean functions they compute by manipulating environmental inputs, enabling a clear distinction between encoded information (genome) and its dynamic execution (epigenome), while allowing precise control of environmental inputs and genetic perturbations [11]. Although Avida does not explicitly implement canonical epigenetic mechanisms, these environmentally driven changes in genome execution provide a functional analogue of epigenetic regulation [12,13].

**Figure 1.**
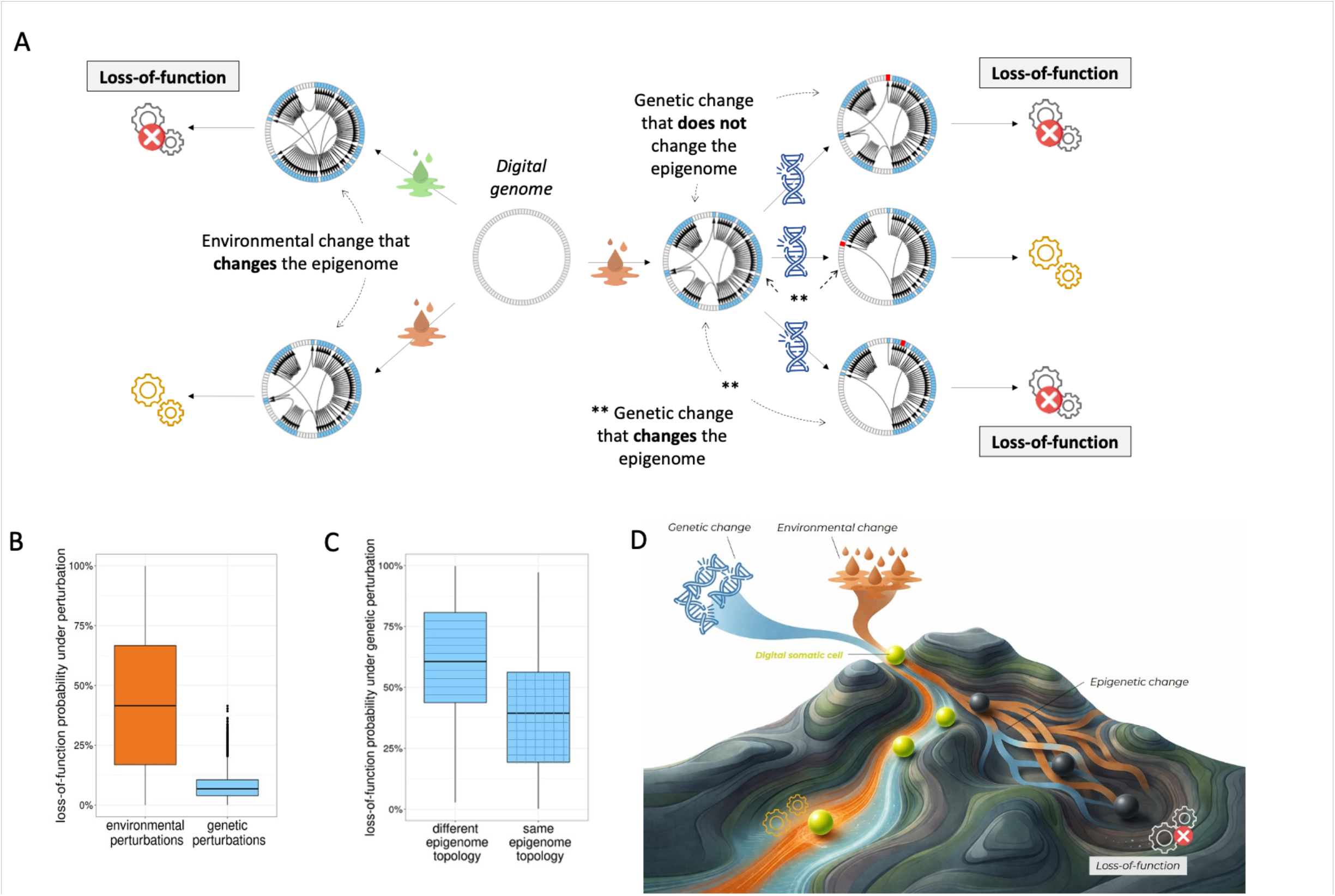
**(A)** Functional loss can arise in digital somatic cells by environmental perturbations (left; droplets) and genetic perturbations (right; single-point mutations, affecting one of the 100 possible genomic positions (mutated positions are indicated in red). The central circular genome (length L = 100 instructions) represents the genetic sequence shared across environments. Blue genomic positions are the ones executed in each epigenome. Colored gears denote functional phenotypes, whereas grey gears indicate non-functional phenotypes. **(B)** Probability of producing a non-functional phenotype across 6,862 genomes after environmental changes (orange) or genetic changes (blue). **(C)** Comparison between changes that reshape the epigenomic topology and changes that preserve the wild-type epigenome. **(D)** Adaptation of Waddington’s epigenetic landscape illustrating how genetic and environmental perturbations reconfigure the epigenome and drive loss of function. Digital cells move across the landscape and settle into functional (yellow) or non-functional (gray) attractors, with environmental (orange) or genetic (blue) perturbations inducing transitions between these states through epigenomic reorganization.

We retrieve 8,177 digital cell genomes (L=100 instructions in length) from avidaDB publicly available database [14] that, when viable (i.e., capable of self-replication), execute at least two distinct epigenomes across 2,500 different environments---one encoding a functional phenotype and one encoding a non-functional phenotype (i.e., losing the ability to compute any Boolean function). Next, we generate all single-point mutants for each genome by replacing the instruction at each position with every other instruction from the genetic language (an alphabet, A, of 26 instructions), resulting in L×(A−1)=2500 mutants per genome. For each somatic mutant, we characterize the resulting epigenomes and phenotypes in a single, common environment; the one that produced the most frequent epigenome encoding a functional phenotype. After excluding genomes whose mutants failed to produce a non-functional phenotype, the final dataset comprised 6,862 genomes. Finally, we compute the probability of transition from a functional epigenome to a non-functional epigenome through both environmental and genetic changes (**Figure 1A**, see **Methods** for more details).

Digital somatic cells that encode functional phenotypes very rarely become cells encoding non-functional phenotypes (1% of cases through environmental changes and 5% through genetic mutations). Nevertheless, environmental or genetic changes do not always result in viable cells. After excluding changes leading to non-viable cells (87% and 26% on average for environmental and genetic changes, respectively), both environmental and genetic changes are still more likely to encode functional than non-functional phenotypes. This tendency is substantially stronger for genetically-induced (log-odds=2.7, p<0.001) than enviromentally-induced changes (log-odds=0.5, p<0.001).

Yet, not all genomic mutated positions are executed. Some occur in inactive regions, analogous to heterochromatin (30% of the genome on average), while others occur in active regions, analogous to euchromatin. Since mutations in inactive regions do not modify the epigenome, they do not change the phenotype. Therefore, after excluding mutations in inactive regions, the likelihood of genetic changesto produce non-functional phenotypes decreased by up to 11% (log-odds=2.1; p<0.001, **Figure 1B**). Moreover, if a genetic mutation occurs in the active region of the genome and results in a non-functional phenotype (i.e., the 11% mentioned above), it is more likely that this phenotype arises through epigenome modification (67%) than without it (33%) (log-odds=0.7; p<0.001, **Figure 1C**). In other words, a significant proportion of genetic mutations that lead to loss-of-function phenotypes do so by altering the epigenome, highlighting the role of epigenomic changes in driving loss of function.

These results indicate that loss-of-function outcomes depend on how perturbations reorganize the epigenomic topology (**Figure 1D**), motivating a quantitative analysis of the determinants of epigenome functionality. We quantified the transition entropy---a metric that quantifies the uncertainty of the next genomic position executed given the current one---as a proxy for epigenome complexity, alongside the number of instructions in the euchromatin (epigenome size) and the richness of their executed instructions (epigenome diversity). A generalized linear mixed-effects model (GLMM) with random intercepts and slopes for genome identity showed that epigenome functionality is strongly influenced by entropy, epigenome size, and richness (**Figure 2**).

**Figure 2.**
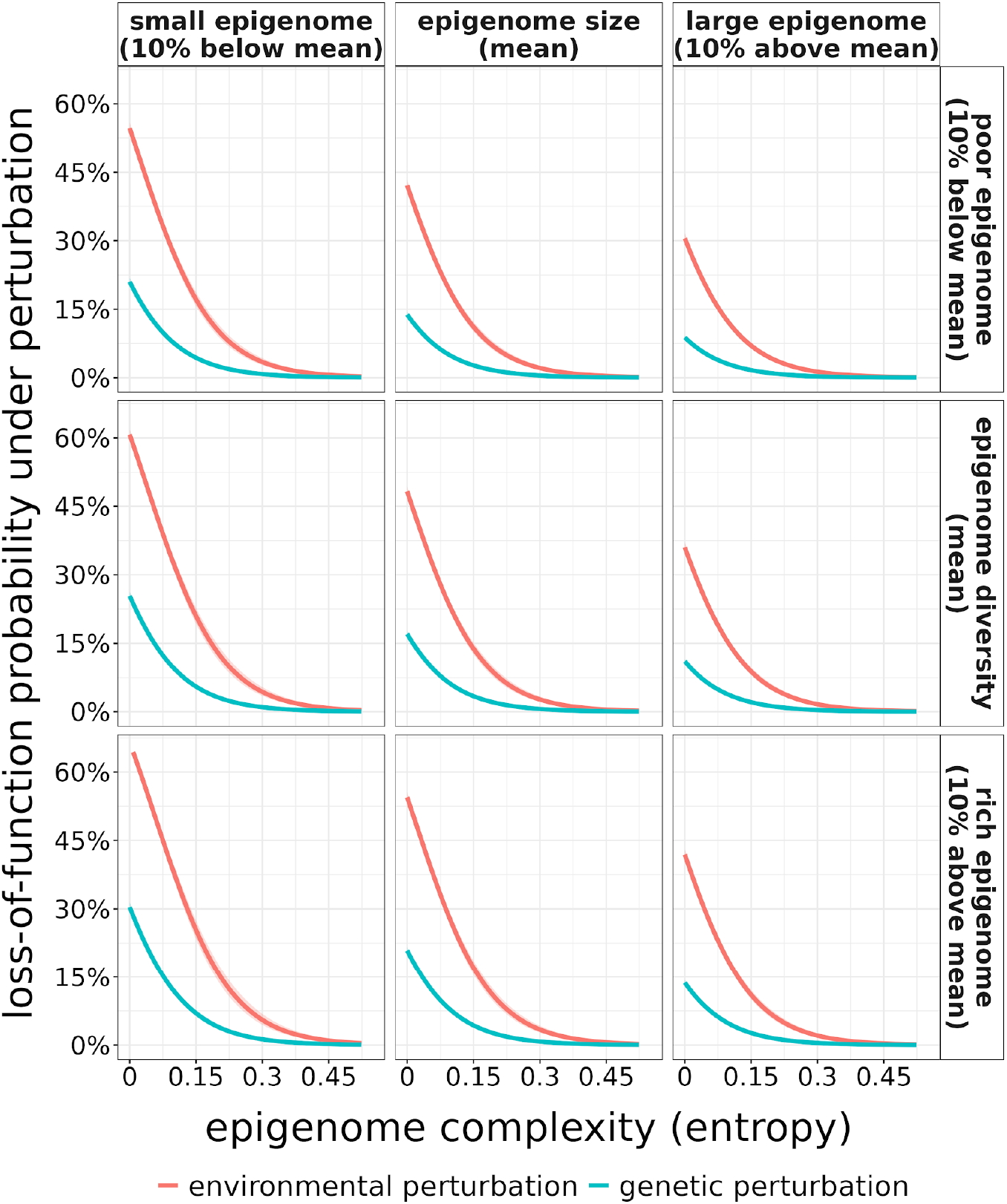
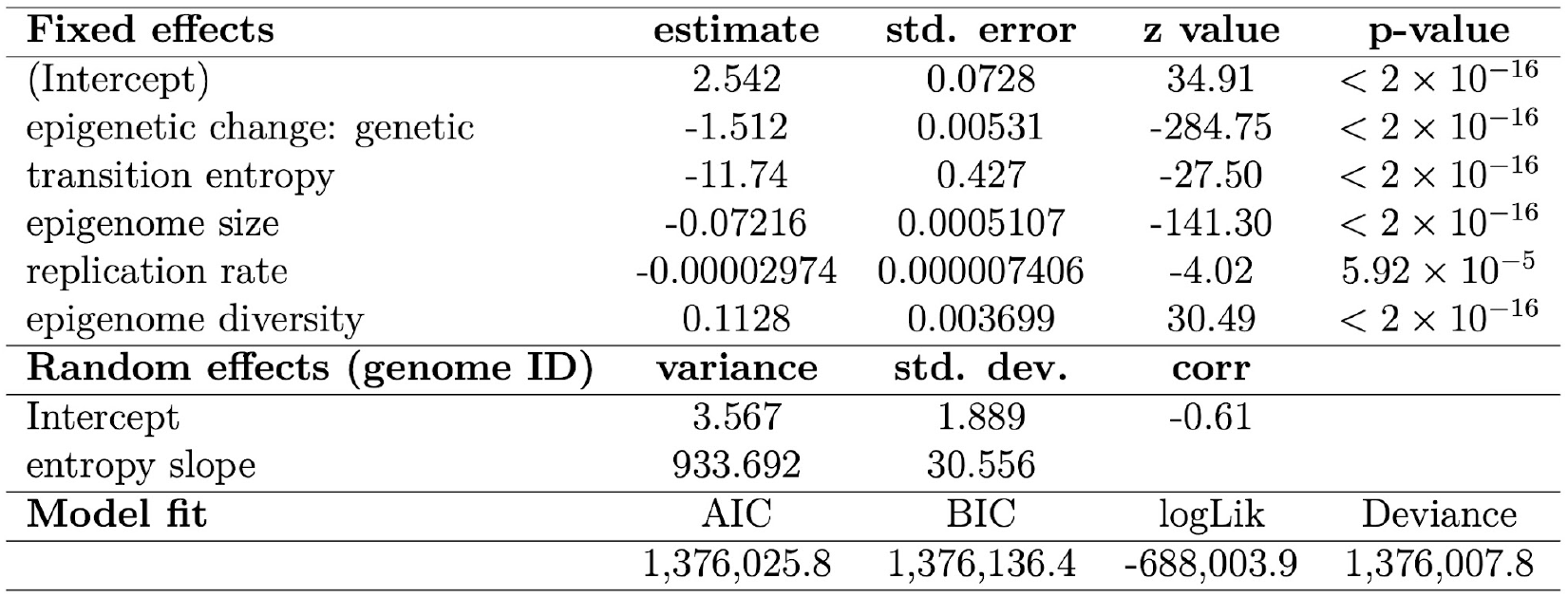
Top: Loss-of-function probability under environmental (red) and genetic (blue) perturbations across epigenomic architectures, as a function of epigenome complexity (transition entropy), shown across combinations of epigenome size and diversity (in sequence instructions). Bottom: GLMM results quantifying these effects.

Increasing entropy by 0.01 reduced the odds of encoding a non functional phenotype by 11%, indicating that slightly higher entropy favors functional outcomes. Each additional instruction in the euchromatin decreased the probability of encoding a non-functional phenotype by 7%. In contrast, each novel instruction executed by the epigenome increased the probability of enconding a non-functional phenotype by 11%. Genomes of digital somatic cells with higher baseline risk were less responsive to changes in entropy, whereas genomes with lower baseline risk were more strongly affected.

Epigenomes encoding functional phenotypes display more unpredictable execution flow than non-functional epigenomes, both under environmental and genetic changes, with average increases of approximately 4% and 11%, respectively. Moreover, the execution flow of epigenomes encoding non-functional phenotypes is 9% more unpredictable when these phenotypes arise from environmental changes than from genetic changes.

Epigenomes that execute a larger fraction of the genome are more likely to encode functional phenotypes, likely because a greater portion of the genome contributes to productive activity. In contrast, epigenomes with a higher number of distinct instruction types are more prone to encode non-functional phenotypes under perturbations, suggesting that while broader genomic accessibility supports functionality, excessive diversity in instruction types can introduce conflicting interactions or errors. Similarly, epigenomes with slightly higher transition entropy—reflecting more complex spatio-temporal patterns of instruction execution—tend to favor functional outcomes.

## Conclusions

Our findings highlight a delicate balance: functional robustness benefits from wide genomic engagement and structured complexity, but too much diversity of potentially interfering instructions or poorly coordinated execution can compromise function. This high transition entropy of functional epigenomes in digital cells is consistent with recent findings that functional regulatory regions exhibit greater stochasticity and more frequent state transitions [15]. Indeed, our results align with models in which environmental fluctuations amplify regulatory noise, whereas mutations tend to produce more predictable epigenomic responses [16,17].

Overall, our results allow us to reinterpret loss of functionality not as cumulative damage, but as a state transition mediated by epigenomic reorganization [1] across a dynamic landscape (**Figure 1D**). Sharing a structural analogy with Waddington’s epigenetic landscape and its recent formalization [18], functional and non-functional phenotypes correspond to alternative attractors within an execution space. Environmental perturbations and mutations do not cause dysfunction directly; rather, they reshape the landscape geometry—modifying attractor stability, altering transition barriers, and driving the system across bifurcation thresholds into non-functional states. Vulnerability depends not on mutational burden alone, but on the system’s propensity to reconfigure its epigenomic architecture.

## Methods

### Digital somatic cells as an experimental system

In Avida, the most widely used computational platform for studying evolution, a digital organism, here interpreted as a somatic cell, is a self-replicating computer program that mutates and evolves within a user-defined computational environment. It consists of a list of instruction codes—its genome—and a virtual CPU that continuously executes these instructions to copy its genome into a new memory space. This process is the only means by which a digital cell can transmit its genetic information to future generations.

### The genome of a digital cell

The genome of a digital somatic cell is a circular sequence in which each site contains one instruction from a 26-instruction programming language, analogous to the four-nucleotide genetic alphabet of carbon-based life. This programming language contains instructions for copying an cells’s genome as well as for storing and manipulating 32-bit binary numbers in buffers, stacks, and registers (AX, BX, and CX).

During replication, the virtual CPU executes instructions according to its genomic sequence, but not all instructions are necessarily executed. Some instructions may never be executed if the instruction pointer—the pointer that determines the next instruction to execute—never reaches their position (digital analogue to histones, which render certain regions of somatic cell chromosomes inaccessible). This can happen because certain instructions modify the execution flow.

For example, some instructions conditionally skip the next instruction depending on the contents of the BX and CX registers (if-less skips if BX > CX; if-n-equ skips if BX = CX). Others move the instruction pointer through the cell’s memory by an amount determined by the CX register (jump-head) or to the position in memory of the flow-head (move-head). Finally, the instruction set-flow moves the flow-head—which marks the start of a loop and allows the instruction pointer to jump back to it—to the memory position specified by CX.

### The epigenome of a digital cell

The epigenome of a digital cell is defined by the positions and order of the genomic instructions that are executed during successful replication. It depends on the content of the BX and CX registers, and its length corresponds to the number of instructions executed during genome replication, among all those encoded in the genome (i.e., the epigenome length is always less than or equal to the genome length).

In contrast, the transcriptome length (the digital analogue of gene expression) is defined as the total number of instruction executions required to complete replication, regardless of their positions or order. The transcriptome length is always greater than or equal to the genome length and is capped at 3,000 instructions, which we set as the threshold for successful replication. Digital cells exceeding this limit are assumed to be unable to replicate.

### The phenotype of a digital cell

The execution of the instructions comprising the epigenome enables a digital cell to compute one or more Boolean functions that characterize its phenotype during replication. This definition arises from the structure of the underlying programming language, which includes instructions capable of storing and manipulating three 32-bit binary numbers in buffers, stacks, and registers (hereafter referred to as the environment of binary inputs processed by the cell) and of generating 32-bit binary outputs.

For example, a Boolean function implementing the logical NAND operation on two 32-bit input numbers outputs a 32-bit value in which each bit is 0 if and only if the corresponding bits of both input numbers are 1, and 1 otherwise. In logical terms, the NAND operation corresponds to the negation of the conjunction (AND) of the two input values. We define a functional phenotype as one that computes at least one Boolean function within a given environment.

### Non-functional phenotypes through environmental changes

The phenotype encoded by the epigenome of a digital cell can be modified by the environment each time an input-output instruction is executed. This instruction reads the 32-bit binary number stored in the cell’s BX register and outputs it, while also checking for any Boolean function performed on the two 32-bit binary numbers stored in its buffers. As a result, the cell’s phenotype depends on the content of the BX register at the moment an input-output instruction is executed.

The content of the BX register can be modified by specific instructions in the cell’s programming language, executed as part of its genome. These include: performing a bitwise NAND on BX and CX and storing the result in BX (nand); adding to or subtracting from BX the content of CX (add, sub); incrementing or decrementing BX by one (inc, dec); swapping BX with CX (swap); moving numbers between BX and the active stack, including toggling the active stack (pop, push, swap-stk); and shifting all bits in BX left or right by one (shift-l, shift-r).

A given environment is defined by a unique set of three 32-bit binary numbers provided to cells at birth. Consequently, an digital cell’s genome will always produce the same epigenome in the same environment, since it receives the same 32-bit input values each time. However, the same genome can give rise to distinct epigenomes in different environments, as the epigenome is conditionally influenced by the content of the BX register during execution. Moreover, even the same epigenome can encode different phenotypes across environments, because the BX register at birth may contain different values, leading the input–output instruction to produce 32-bit numbers that correspond to the computation of different Boolean functions.

### Non-functional phenotypes through genetic changes

To reproduce, a digital cell must copy its genome instruction by instruction into a new region of memory, a process that may introduce errors (i.e., mutations). A mutation occurs when an instruction is copied incorrectly and replaced in the offspring genome by another instruction chosen at random, with uniform probability, from the 26-instruction programming language.

Because such mutations can alter the execution flow—and consequently the epigenome executed by the cell—the resulting phenotype may differ from that of the parent. Moreover, as in the case of environmental changes, a mutation may cause a loss of function without altering the epigenome executed by the parent, because it changes the content of the BX register rather than the execution flow.

## Computational study design

### Dataset

We retrieved the subset of 8,177 digital cells with genomes of length L = 100 from avidaDB that, when viable (i.e., capable of self-replication), executed across 2,500 distinct environments at least one epigenome encoding only the non-functional phenotype and at least one encoding one or more functional phenotypes, with no epigenome encoding both.

### Identifying environmental changes

We ran each of the 8,177 target genomes in isolation and quantified, in addition to the epigenomes they executed and the phenotypes they encoded, their viability across the 2,500 environments (i.e., the fraction of environments in which the digital cell was capable of replication).

For each distinct epigenome, we recorded whether it encoded the non-functional phenotype, any functional phenotypes, the number of environments in which it occurred, its length, and its transcriptome length. The likelihood of encoding the non-functional phenotype through environmental changes was quantified for each genome as the fraction of its viable epigenomes that encode the non-functional phenotype.

In addition to the target dataset—genomes executing epigenomes that encode either non-functional or functional phenotypes, but not both—we analyzed the likelihood of encoding the non-functional phenotype in other genome groups, classified according to the phenotypes encoded by their epigenomes:

- Genomes executing epigenomes that encode only non-functional phenotypes, only functional phenotypes, or both.
- Genomes executing epigenomes that encode only non-functional phenotypes and both non-functional and functional phenotypes.
- Genomes executing epigenomes that encode only functional phenotypes and both non-functional and functional phenotypes.
- Genomes executing only epigenomes that encode both non-functional and functional phenotypes.

### Identifying genetic changes

For each of the 8,177 targeted genomes, we generated all L × (A − 1) = 2,500 possible single-point mutants, where L denotes the genome length and A the size of the instruction alphabet. Then, each target genome’s mutants were run in one of the environments—identical for all mutants—in which the target genome executed its most frequent epigenome encoding a functional phenotype.

For each mutant, we recorded the resulting epigenome, the phenotypes it encoded (one or more per epigenome), its viability, and both its length and transcriptome length. We then excluded target genomes whose mutants did not produce a non-functional phenotype, resulting in a final set of 6,862 genomes whose mutants, at least one, executed both the wild-type epigenome and at least one epigenome encoding the non-functional phenotype.

The likelihood of encoding the non-functional phenotype through genetic changes was quantified for each genome as the fraction of viable epigenomes, executed by its mutants, that encode the non-functional phenotype.

## Declarations

### Availability of data and materials

All data and code are available at Zenodo (DOI: 10.5281/zenodo.21414319) [19].

### Author’s contributions

MAF mined AvidaDB data and performed all the *in silico* experiments. All authors designed research, analysed data, prepared figures and wrote the paper.

